# Characterizing Trabecular Bone Properties near the Glenohumeral Joint Following Brachial Plexus Birth Injury

**DOI:** 10.1101/2020.02.26.967224

**Authors:** Emily B. Fawcett, Carolyn M. McCormick, Austin F. Murray, Dustin L. Crouch, Katherine R. Saul, Jacqueline H. Cole

## Abstract

Brachial plexus birth injury (BPBI) causes functional arm impairment in 30-40% of those affected due to altered loading on the glenohumeral joint. While gross morphological osseous deformities have been seen in the humerus and scapula, alterations in the underlying trabecular bone microstructure and mineralization are not clear. Using a murine model of BPBI, trabecular bone alterations were explored in the proximal humerus and distal scapula, which surround the articulating surface of the joint. Samples were scanned using micro-CT, reoriented, and analyzed for standard trabecular metrics. The regions of interest closest to the articulating surface showed the greatest detriments. In the scapula, the scapular neck region showed less robust trabecular bone in the neurectomy group with decreased BV/TV (p=0.001), BMD (p=0.001), Conn.D (p=0.006), Tb.N (p<0.0001), and DA (p=0.033), and increased Tb.Sp (p<0.0001) compared to sham. In the humerus, the epiphysis showed less robust trabecular bone in neurectomy group, but to a much lesser extent than the scapular neck. The neurectomy group showed reduced BMD (p=0.007) and Tb.N (p=0.029) compared to sham. Data suggest deformities are worse near the articulating surface, likely due to the greater amount of mechanical loading. The reduction in trabecular microstructure and mineralization may compromise bone strength of the affected limb following BPBI. Further investigation of the underlying trabecular bone deformities following injury are necessary to eventually inform better treatments to limit the development of deformities.

## Introduction

Functional impairment of the arm is permanent in 30-40% of children who sustain brachial plexus birth injuries (BPBI) during difficult childbirth (Pondaag, 2004).^(1)^ Secondary glenohumeral dysplasia in the affected limb can result from injury (Waters 1998) and may alter the mechanical loading of the joint, which is critical for normal bone and joint development. Regions in the glenohumeral joint experience abnormal growth and macrostructural deformities after BPBI^(2–6)^ (Al-Qattan 2003, Nikolaou 2011, Crouch 2014, Crouch 2015, Cheng 2015), and are comprised primarily of cancellous bone, which plays an integral role in transferring joint loading along the bone and is known to adapt to altered loads (Wolff, 1892/1986; Frost, 1994).

Trabecular bone loss has been observed in many reduced loading scenarios, such as spaceflight^(7)^ (Vico *et al.*, 2000), prolonged bedrest^(8)^ (Kazakia *et al.*, 2014), and spinal cord injury (Eser *et al.*, 2004).^(9)^ However, the nature and extent of trabecular bone changes after BPBI are not well understood. While no human BPBI study has examined bone microstructural properties, several studies using murine BPBI models have reported trabecular bone detriments in the humeral head, with fewer and thinner trabeculae and greater separation compared to contralateral limb and sham control groups^(10–12)^ (Kim, 2009; Kim, 2010; (Potter, 2014). These trabecular changes were observed in conjunction with global musculoskeletal changes, including humeral anteversion and glenoid retroversion^(10,11)^ (Kim *et al.*, 2009; Kim *et al.*, 2010), as well as decreased supraspinatus muscle volume^(10–12)^ (Kim *et al.* 2009; Kim *et al.*, 2010; Potter *et al.*, 2014). Beyond morphological changes, the elastic modulus and overall strength of the humeral head decreased (Potter 2014).^(12)^ However, previous animal studies have not characterized trabecular changes throughout the entire glenohumeral region, particularly in the scapula glenoid region and humeral metaphysis, nor have they examined the relationship between changes in glenohumeral macrostructure and changes in the underlying trabecular microstructure. In addition, tissue mineral density is related to elastic modulus, which has been seen to decrease. Yet tissue mineral density has not been characterized following BPBI. This information could reveal regions of the glenohumeral joint that may be especially susceptible to BPBI-induced changes in trabecular bone, ultimately informing better treatment strategies to mitigate the macrostructural deformities and address the deficits in arm function experienced by human patients.

Mechanical loading is well established as a critical determinant of bone morphology throughout development (Frost, 1994; Turner, 1998).^(13,14)^ Altered mechanical loading has been proposed as a major contributing factor in the development of osseous deformities in the glenohumeral joint affected by BPBI. Osseous deformities reported in clinical studies include flattening of the humeral and posterior subluxation relative to the scapula (Reading 2012, Waters 1998).^(15,16)^ Most scapular deformities have been identified in the glenoid fossa, including glenoid retroversion, declination, loss of glenoid concavity, and pseudoglenoid formation (Waters 1998).^(16)^ Computational musculoskeletal models developed from animal studies have determined that, when compared to a normal developing shoulder, the BPBI glenohumeral joint experiences increased posteriorly directed, compressive loads, due to impaired growth of surrounding muscles (Crouch, 2014, Cheng, 2015).^(4,6)^ Indications of altered mechanical loading following BPBI motivate further investigation of the glenohumeral trabecular microstructure and mineralization, which bears primary responsibility for distributing loads in the shoulder joint.

The purpose of this study was to characterize microstructural features in the underlying trabecular bone near the articulating glenohumeral joint (e.g., humeral epiphysis and metaphysis, scapular regions near the glenoid) using an established rat model of BPBI (Li *et al.*, 2008; Nikolaou *et al.*, 2015).^(17,18)^ Given the supporting evidence from other scenarios of altered loading and the remarkable skeletal dysplasia known to occur in clinical cases of BPBI, we hypothesized that trabecular tissue mineral density and microstructure in the affected shoulder are compromised during postnatal development following BPBI.

## Methods

### Study Design

This study was performed using humeri and scapulae obtained from a previous study (Crouch 2015).^(19)^ All animal procedures were approved by the Institutional Animal Care and Use Committee at the Wake Forest School of Medicine. Sixteen Sprague Dawley rat pups (Harlan Laboratories, Indianapolis, Indiana) were grouped according to surgical intervention implemented five days after birth: neurectomy and sham (*n*=8 each) (Figure 1).

**Figure 1.**
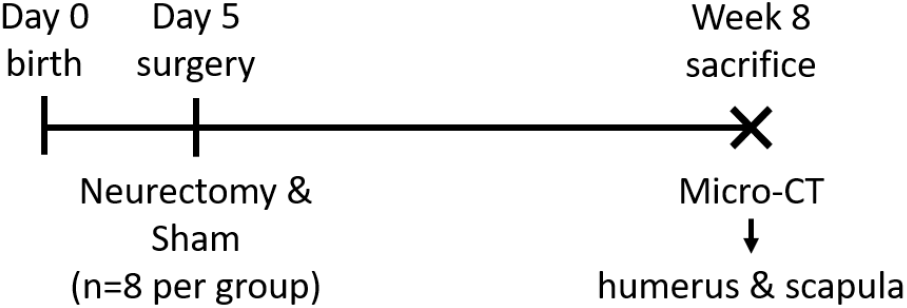
Overview of study timeline. Interventions were performed at the same timepoint. Analyses will be performed following sacrifice at week 8.

In the neurectomy group, a postganglionic injury was surgically induced in the left forelimb using an established model (Li 2008),^(17)^ and the right forelimb was kept completely intact as a control. The rats were anesthetized with inhaled isoflurane, and a small transverse infraclavicular incision was made, splitting the left pectoralis major muscle to expose the brachial plexus. The left C5 and C6 nerve roots and the upper trunk of the brachial plexus were transected distal to the dorsal root ganglion. The wound was irrigated with saline and closed with 6-0 nylon suture. The sham group received a similar surgery with incision through the left pectoralis major muscle, but the brachial plexus was not transected, and the right forelimb was kept completely intact.

After eight weeks, the rats were sacrificed, and the left (affected) and right (unaffected) scapulae and humeri were harvested. The bones were fixed in 10% neutral buffered formalin for 48 hours and then immersed in 70% ethanol for storage. Two scapulae per group and one humerus per group were damaged during the original dissection, leaving n = 6 left and right scapulae and n = 7 left and right humeri for this study.

### Micro-Computed Tomography

Trabecular bone density and microstructure were assessed with quantitative micro-computed tomography (micro-CT). The bones were scanned in 70% ethanol using a 0.5-mm Al filter, 45 kVp and 177 μA, 800-ms integration time, 1,000 projections per rotation, and no frame averaging (μCT 80, SCANCO Medical AG, Brüttisellen, Switzerland). Density measurements were calibrated using the SCANCO hydroxyapatite calibration phantom, and a threshold of 441 mg/cm^3^ (3891 Hounsfield units) was applied for bone. The scans were reconstructed at an isotropic voxel size of 10 μm and reoriented for consistent anatomical alignment. Trabecular bone volumes of interest (VOIs) were selected near the glenohumeral joint, which was expected to be most affected following nerve injury. Three VOIs in the glenoid fossa region of the scapula and two VOIs in the proximal humerus were selected by manually contouring the trabecular bone in a series of slices and then using automated morphing interpolation across these contours to create the volumes (Figure 2). All VOI lengths were chosen based on the maximum possible length in each region for the smallest bone.

**Figure 2.**
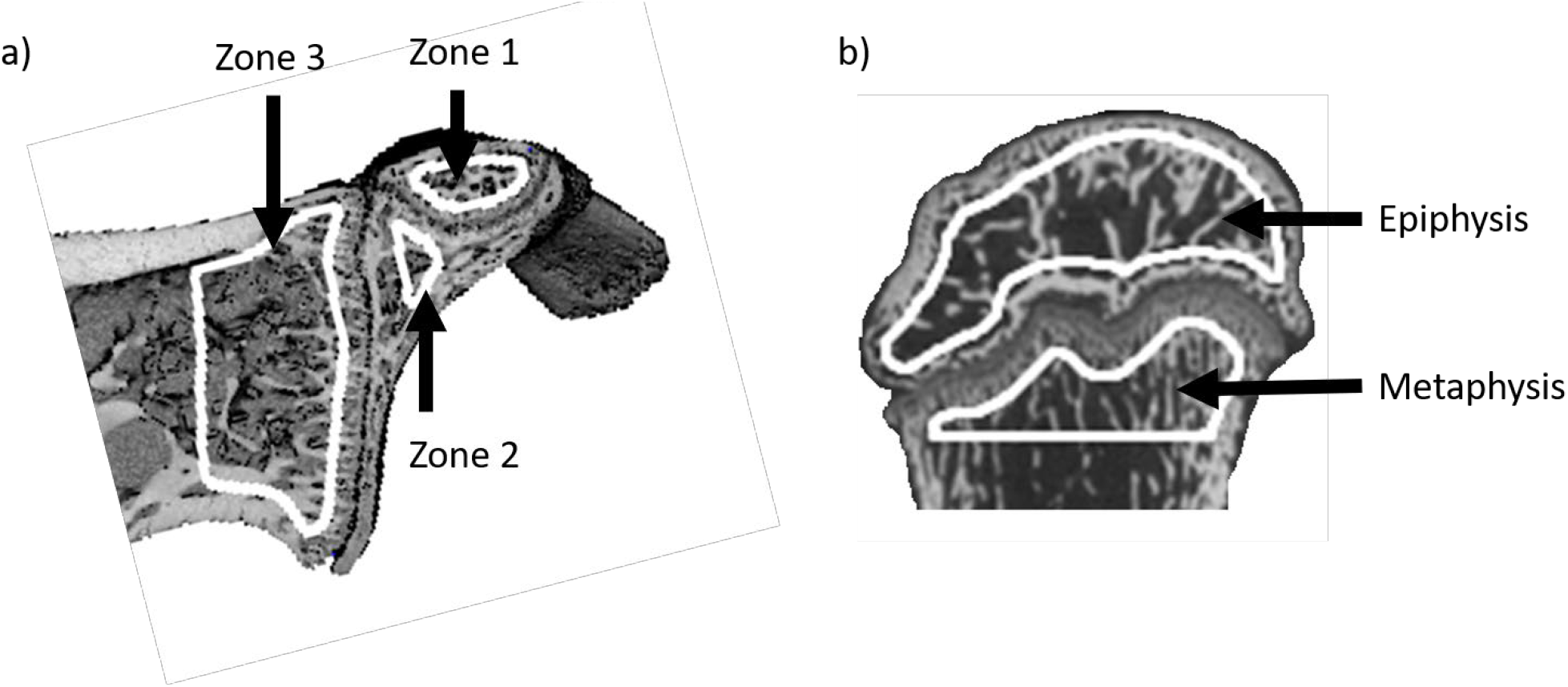
Trabecular bone volumes of interest (outlined in white) in the a) scapula and b) humerus, selected within different ossification centers, or zones.

The three scapular VOIs were defined in secondary ossification centers formed during postnatal development (Kothary *et al.*, 2014).^(20)^ The first scapular VOI (*zone 1*) was 6.5% of the total scapular length, defined as the distance from the beginning of the glenoid region to the farthest point of the proximal end. This VOI was positioned in the subcoracoid secondary ossification center, beginning next to the articular surface and extending proximally along the scapular spine axis toward the superior glenoid physis. The second scapular VOI (*zone 2*) was 1.5% of the total scapular length and defined in the upper glenoid secondary ossification center, beginning inferior to the superior glenoid physis and next to the articular surface extending proximally along the scapular spine. The third VOI (*zone 3*) was 7.5% of the total scapular length and was defined within the scapular neck, beginning next to the physis running across the width of the neck and extending proximally. The first humeral VOI was 12.5% of the total humeral length and defined in the epiphysis, beginning inferior to the articular surface and extending distally toward the proximal growth plate. The second humeral VOI was 5% of the total humeral length and defined in the metaphysis, beginning inferior to the proximal growth plate and extending distally toward the diaphysis. For each VOI, standard trabecular bone metrics were calculated using the SCANCO analysis software, including bone volume fraction (BV/TV), bone mineral density (BMD), tissue mineral density (TMD), connectivity density (Conn.D), trabecular number (Tb.N), mean trabecular thickness (Tb.Th) and separation (Tb.Sp) using direct 3D methods (Hildebrand and Rüegsegger 1997),^(21)^ and degree of anisotropy (DA) (Bouxsein *et al.*, 2010).^(22)^

### Statistical Analyses

Four sets of analyses were performed. For analysis 1, limb comparisons for trabecular bone metrics were examined between affected and unaffected limbs within each group (sham, neurectomy) using paired t-tests with Welch’s correction for unequal variances. For analysis 2, group comparisons for the affected-to-unaffected ratios were examined between the neurectomy and sham groups using unpaired t-tests with Welch’s correction for unequal variances. For analysis 3, anatomical site relationships were examined between the scapula and humerus metrics using linear correlations. These relationships were evaluated using both the microstructural metrics from this study and the macrostructural metrics from the previous study using these same bones (humeral head width, thickness, and curvature; glenoid inclination and curvature) (Crouch 2015).^(5)^ For analysis 4, length-scale relationships between the glenohumeral joint macrostructure and underlying trabecular microstructure were assessed by stepwise multiple regression with forward selection, using Schwarz Bayesian information criterion to determine which trabecular microstructural properties (predictor variables) best explained the variability in the macrostructural measurements in the humeral head (width, thickness, and curvature) and glenoid fossa (inclination and curvature). Microstructural values from the humeral epiphysis and glenoid zone 3 (Z3) were used for the correlations and multiple regressions, since these regions were closest to, and thus most relevant to, the articulating glenohumeral joint. Analyses 1, 2, and 3 were completed with Prism 6 (GraphPad Software, Inc., La Jolla, CA), and analysis 4 was completed with SAS 9.4 (SAS Institute Inc., Cary, NC). Significance was defined as p < 0.05, and trends were defined as p < 0.08.

## Results

### Limb Comparisons

#### Scapula

For the neurectomy group, trabecular bone density and microstructure were altered in the scapula of the affected limb when compared to the contralateral unaffected limb (Table 1, Figure 3). Bone volume fraction was decreased 21-26% in zones 2 and 3 and tended to decreased about 18% in zone 1, and bone mineral density was decreased 19-24% in zones 1, 2, and 3, indicating a reduction in bone mass on the affected side. Zone 2 also tended to show decreased tissue mineral density, indicative of material change. In zones 1 and 3, trabecular number was reduced by approximately 20% and trabecular separation was increased 35-44% on the affected side. Zone 2 had decreased trabecular thickness by approximately 18%. The alterations in trabecular architecture indicate less robust trabecular bone in zones 1 and 3 on the affected side compared to zone 2. In addition, zone 1 portrayed a decrease in anisotropy of about 10%.

**Table 1.** Trabecular bone metrics in the scapula volumes of interest (VOI), expressed as percent difference between the affected and the unaffected side. ^#^p < 0.1, *p < 0.05, **p < 0.01 for affected vs. unaffected.

| Surgery | VOI | Affected Side vs. Unaffected Side (%) |  |  |  |  |  |  |  |
| --- | --- | --- | --- | --- | --- | --- | --- | --- | --- |
|  |  | BV/TV | BMD | TMD | Conn.D | Tb.N | Tb.Th | Tb.Sp | DA |
| Sham | Zone 1 | 3.9 | 2.6 | -1.2 | -3.1 | 3.5 | 2.9 | -3.4 | -4.3 |
|  | Zone 2 | -5.4** | -5.2** | -2.1 | 22.4 | 7.8 | -3.7 | 12.4 | 30.6 |
|  | Zone 3 | 1.9 | 0.6 | -1.6 | 11.5* | 4.8<sup>#</sup> | -2.8 | -<br>5.0<sup>#</sup> | 3.6 |
| Neurectomy | Zone 1 | -17.8<sup>#</sup> | -19.4* | -3.9 | -15.2 | -18.8* | 0.9 | 44.0** | -9.7* |
|  | Zone 2 | -25.6** | -23.9** | -5.3<sup>#</sup> | -4.3 | -3.0 | -18.3** | 15.6 | 5.6 |
|  | Zone 3 | -20.5* | -21.8* | -3.5 | -18.2<sup>#</sup> | -20.1* | -2.6 | 34.8* | -10.2 |

**Figure 3.**
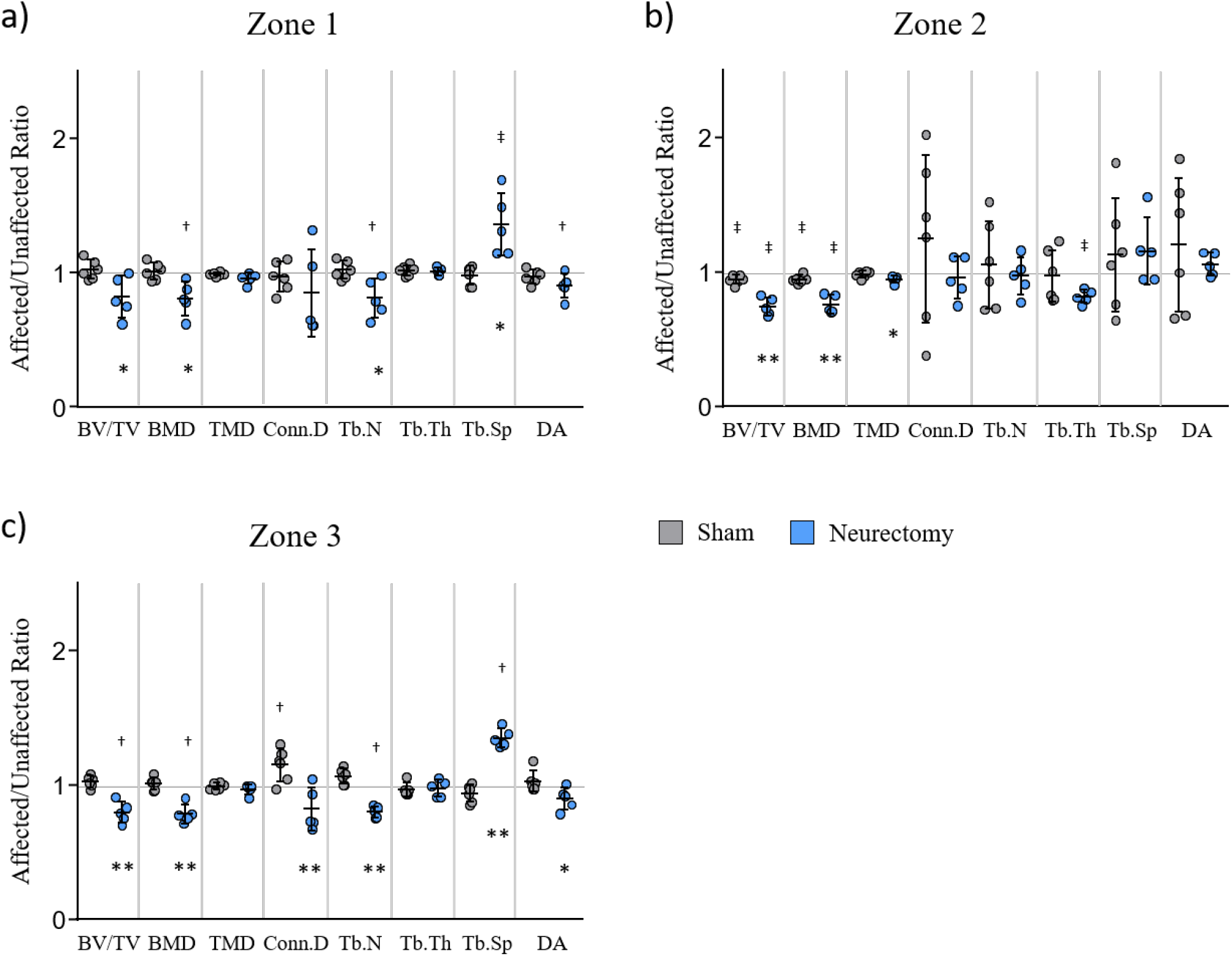
Trabecular bone metrics in the scapula. All three zones exhibited differences in trabecular microstructure in the neurectomy group with zone 3 portraying the most differences. a) Zone 1 shows decreased BV/TV, BMD, and Tb.N, and increased Tb.Sp. b) Zone 2 shows decreased BV/TV, BMD, and TMD. c) Zone 3 shows decreased BV/TV, BMD, Conn.D, Tb.N, and DA, and increased Tb.Sp. ^†^p < 0.05, ^‡^p < 0.01 affected vs. unaffected limb, *p < 0.05, **p < 0.01 neurectomy vs. sham.

The sham group also experienced some differences in trabecular bone microstructure for zones 2 and 3 of the scapula on the affected side when compared to the contralateral limb (Table 1, Figure 3). Specifically, zone 2 had reduced bone volume fraction and bone mineral density by about 5%, indicating decreased bone mass. In addition, zone 3 had a 12% increase in connectivity density and tended to have a 5% increase in trabecular number and 5% decrease in trabecular separation. However, the bone on the affected sides in the sham group were much less affected than those in the neurectomy group.

#### Humerus

The neurectomy group exhibited less robust trabecular bone in the affected epiphysis when compared to the unaffected epiphysis (Table 2, Figure 4a). Bone volume fraction and bone mineral density were reduced by approximately 19% and 24% respectively, revealing loss of bone mass in the affected limb. Related to material changes, tissue mineral density tended to decrease by 4% on the affected side. In addition, trabecular separation was increased by about 90% on the affected side and trabecular number tended to decrease by 33%, indicating less robust trabecular architecture. In the metaphysis, less alterations were present in the affected limb (Table 2, Figure 4b). Trabecular material was significantly altered with a 6% reduction in tissue mineral density, and connectivity density was increased by approximately 112%.

**Table 2.** Trabecular bone metrics in the humerus volumes of interest (VOI), expressed as percent difference between the affected and the unaffected side. ^#^p < 0.1, *p < 0.05, **p < 0.01 for affected vs. unaffected.

|  |  | Affected Side vs Unaffected Side (%) |  |  |  |  |  |  |  |
| --- | --- | --- | --- | --- | --- | --- | --- | --- | --- |
| Surgery | VOI | BV/TV | BMD | TMD | Conn.D | Tb.N | Tb.Th | Tb.Sp | DA |
| Sham | Epiphysis | 0.5 | 1.2 | -0.9 | 0.5 | 0.2 | 0.1 | -0.9 | 0.4 |
|  | Metaphysis | 14.9 | 18.5 | -2.5 | 139.2** | 16.3* | -6.2<sup>#</sup> | -18.3** | 44.9* |
| Neurectomy | Epiphysis | -19.1* | -24.3** | -3.6<sup>#</sup> | -39.2 | -33.4<sup>#</sup> | 17.6 | 90.1** | -17.2 |
|  | Metaphysis | -8.1 | -6.8 | -5.7* | 112.0* | 16.9 | -14.0 | -10.9 | 21.8 |

**Figure 4.**
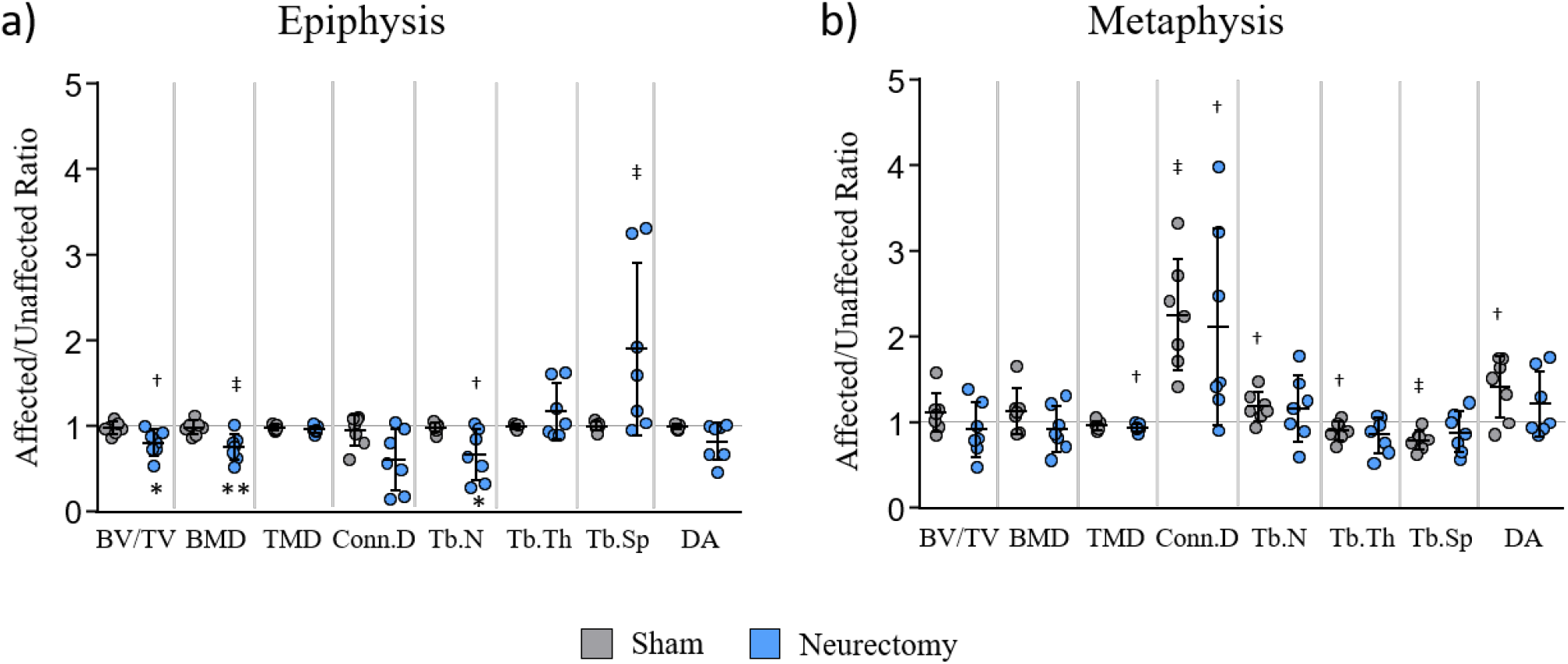
Trabecular bone metrics in the humerus. a) The epiphysis showed decreased BMD, Conn.D, and Tb.N in the neurectomy group. b) The metaphysis showed no significant differences between groups. ^†^p < 0.05, ^‡^p < 0.01 affected vs. unaffected limb, *p < 0.05, **p < 0.01 neurectomy vs. sham.

Unlike in the neurectomy group, in the sham group, there were no significant deformities found in the epiphysis of the affected side when compared to the contralateral limb. However, the metaphysis region of the affected side portrayed increased connectivity density by 139%, trabecular number by 16%, and degree of anisotropy by 45%, and decreased trabecular separation by 18%. Trabecular thickness also tended to decrease by 6%, all indicating different trabecular architecture between limbs.

### Group Comparisons

#### Scapula

In zone 1 and 2 of the glenoid fossa region, few trabecular metrics differed significantly between the neurectomy and sham groups (Figure 4a-b). Zone 1 showed lower bone volume fraction (−19.6%, p=0.039), bone mineral density (−19.8%, p=0.018), and trabecular number (−20.6%, p=0.028), and increased trabecular separation (+38.8%, p=0.020) when compared to sham. Zone 2 showed lower bone volume fraction (−21.3%, p=0.001), bone mineral density (−20.0%, p=0.002), and tissue mineral density (−3.1%, p=0.047) compared to sham. Glenoid trabecular bone was most altered in zone 3 following neurectomy (Figure 4c). Bone volume fraction and bone mineral density were reduced (−21.2%, p=0.001; −22.8%, p=0.001 respectively) compared to sham, representing decreased bone mass following injury. Trabecular architecture was also less robust, indicated by a decrease in connectivity density (−28.7%, p=0.006), trabecular number (−24.5%, p<0.0001), and degree of anisotropy (−11.76%, p=0.033), and an increase in trabecular separation (+45.2%, p<0.0001).

#### Humerus

The epiphyseal region shows few altered trabecular metrics in the neurectomy group (Figure 5a). Specifically, there was a decrease in bone volume fraction (+7.76%, p=0.024), bone mineral density (−16.3%, p=0.007), and trabecular number (−0.3%, p=0.029), and a trend towards decreased connectivity density (−34.0%, p=0.051). and increased trabecular separation (+39.4%, p=0.057) when compared to sham. These altered metrics indicate reduction in bone mass and less robust trabecular architecture.

In the metaphyseal region, there were no significant differences between the neurectomy and sham group (Figure 5b). There were also no trends towards less robust trabecular bone in the neurectomy group compared to sham, suggesting no alterations due to injury.

### Anatomical Site Relationships

Correlations were present between microstructural regions near articulating glenohumeral surfaces (Table 3). Some correlations were present between microstructural regions near the articulating glenohumeral surfaces (Table 3). Tissue mineral density in scapular zone 3, representative of material properties, correlated with epiphyseal trabecular bone architecture metrics and tended to correlate with epiphyseal bone mas metrics. Specifically, scapular zone 3 tissue mineral density positively correlated with epiphyseal connectivity density (r=0.913, p=0.030) and trabecular number (r=0.946, p=0.015), and negatively correlated with trabecular separation (r= −0.978, p=0.004). Tissue mineral density also tended to positively correlate with bone volume fraction (r=0.837, p=0.077) and bone mineral density (r=0.816, p=0.092).

**Table 3.**
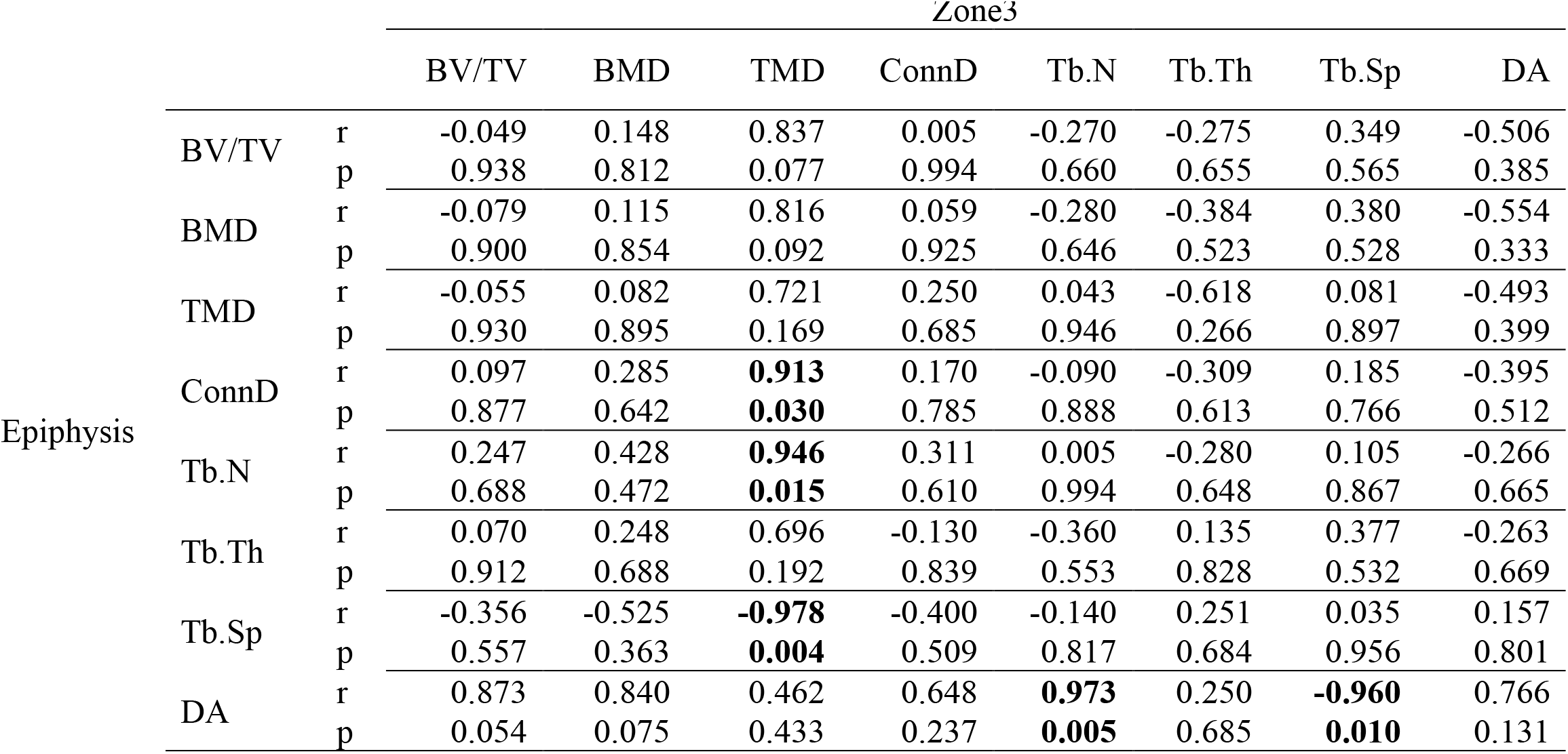
Pearson’s correlation coefficients (r) and associated p-values for correlations of trabecular density and microstructure of the epiphysis and scapular zone 3 in the neurectomy groups. p < 0.05 in bold.

**Table 4.**
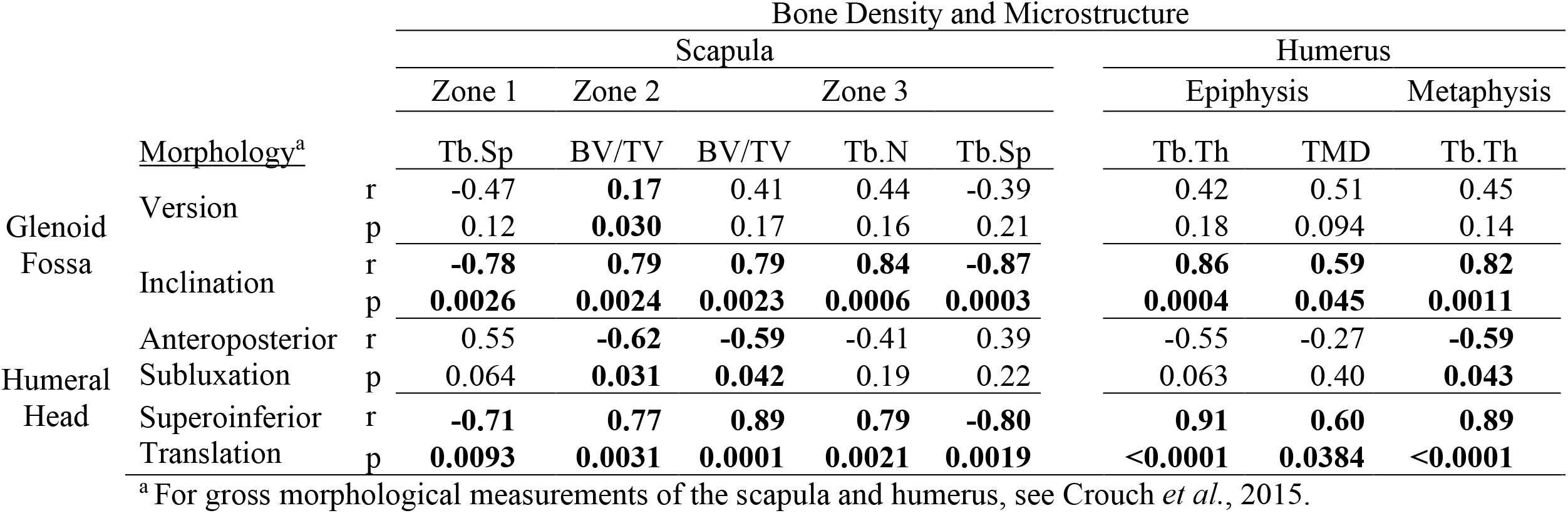
Pearson’s correlation coefficients (r) and associated p-values for correlations of trabecular density and microstructure with macrostructure metrics in BPBI rat shoulders. p<0.05 in bold.

In the epiphysis, degree of anisotropy correlated with some trabecular bone architecture metrics and tended to correlate with bone mass metrics, all from scapular zone 3. Specifically, degree of anisotropy positively correlated with zone 3 trabecular number (r=0.973, p=0.005) and negatively correlated with zone 3 trabecular separation (r= −0.960, p=0.010). The degree of anisotropy in the epiphysis also tended to positively correlate with zone 3 bone volume fraction (r=0.873, p=0.054) and bone mineral density (r=0.84, p=0.075).

### Length-Scale Relationships

Nearly all macrostructural bone measurements (glenoid curvature and inclination, humeral head width, thickness, and curvature) were linearly correlated with at least one microstructural bone metric (Table ##). Multiple regression models included measurements from both groups (sham and neurectomy). Changes in glenoid curvature were best explained by BV/TV and TMD, respectively. Glenoid inclination variability was best explained by Tb.Sp, Conn.D, Tb.N, and SMI. Changes in humeral head thickness were best explained by DA, Tb.Th, TMD, and Tb.Sp. Variation in humeral head curvature was best explained by Conn.D. Humeral head width did not appear to be significantly affected by any particular microstructural parameter.

## Discussion

The goal of this study was to investigate whether trabecular bone density and microstructure are altered in regions near the glenohumeral joint following brachial plexus birth injury in rats. Our findings show clear differences in the trabecular bone density and microstructure for limbs affected by neurectomy compared to both their unaffected contralateral control limb and sham control. The most profound differences in trabecular metrics occurred in the articulating regions of the glenohumeral joint – the humeral epiphysis and the glenoid zone 3. The scapula was affected in more parameters of trabecular bone density and microstructure relative to the humerus. Our findings in the underlying trabecular bone near the glenohumeral joint provide broader context for the global musculoskeletal deformities known to occur in children with BPBI.

Prior work in murine models of BPBI have reported microstructural changes to the humeral epiphysis that worsen over time, but the parameters affected differ from the current report. For example, a cross-sectional study in a CD-1 mouse model of BPBI employing the same type of neurectomy at 24 hours following birth reported a 22.2% reduction in Tb.Th in the humeral epiphysis of the affected limb relative to the contralateral limb and 17.7% relative to sham at four weeks post-injury. After twelve weeks, Tb.Th reduced further to 42.2% relative to the contralateral limb and 50% relative to sham (Kim *et al.*, 2010). ^(13)^ In contrast, we observed reduced epiphyseal BV/TV and BMD, increased Tb.Sp, and a trend towards decreased TMD and Tb.N in the affected neurectomy limb compared to the contralateral control limb and reduced BMD, Conn.D, and Tb.N compared to sham, but no significant changes to Tb.Th. These trabecular bone losses indicate that BPBI compromises bone microstructure shortly after injury and more severely over time, suggesting that timing of post-injury treatment is critical to bone health. These results, combined with the examination of the metaphyseal region and regions of the scapula help reveal the location of the greatest disparities in affected shoulders compared to contralateral or sham shoulders.

Our group is the first to investigate the trabecular microstructure following BPBI in the humeral metaphysis and regions of the scapula within the glenoid fossa. This study found that there was a significant reduction in TMD of the affected neurectomy limb compared to the contralateral limb in both the epiphysis and metaphysis, and a significant reduction in TMD of the neurectomy group in the epiphysiscompared to sham. However, there were no group differences found in the metaphysis. Data suggest deformities are worse near the articulating surface, likely due to the greater amount of mechanical loading. Following analyses of the humerus, this study investigated the effects of BPBI on the scapula. Of the three zones, zone 3 had the least robust metrics. Zone 3 resides closest to the articulating surface, also suggesting greater deformities near the articulating surface. When compared to the humerus, the scapula had greater disparities, especially in the trabecular architecture. The large amount of alterations in scapular microstructure suggest the humerus is not affected the most post-injury.

Although the humerus was less affected than the scapula, some microstructure metrics of the epiphysis did seem to correlate or trend towards a correlation with the scapular zone 3 microstructure metrics. The metrics most correlated were the scapular zone 3 tissue mineral density with epiphyseal trabecular bone architecture metrics and bone mass metrics and the epiphyseal degree of anisotropy with zone 3 trabecular bone architecture metrics and bone mass metrics. This suggests that when the trabecular bone in the humeral epiphysis becomes less robust, the tissue mineral density in the scapular neck is also altered. It also suggests that when the trabecular bone in the scapular neck becomes less robust, the degree of anisotropy in the humeral epiphysis increases, meaning the trabecular bone orientation is more anisotropic. However, the trabecular metrics as a whole are not changing similarly between the two regions. Therefore, the mechanical loading may be affecting one bone more than the other and not to a similar degree.

Based on previous studies of human trabecular bone, even modest decreases in bone tissue mineralization, such as the ~4% drop in TMD in the scapular zone 2, can lead to much greater reductions in the tissue modulus and strength (Carter 1977),^(23)^ which will weaken bone’s overall resistance to loading. Diminished physical properties of trabeculae (e.g., BV/TV, Tb.N, Tb.Th, or Tb.Sp) can also increase fracture risk, as has been shown in other clinical cases like osteoporosis^(24)^ (Parfitt *et al.*, 1983)^(38)^ and osteopenia (Stein *et al.*, 2014).^(25)^ Reduced Tb.N is a significant predictor of vertebral fracture in middle-aged men diagnosed with osteopenia (Legrand *et al.*, 2000).^(26)(40)^ Although it has not been studied, our findings suggest that the alterations in trabecular architecture following BPBI may decrease load-bearing capacity in the affected limb and increase the risk of bone fracture in the humerus and scapula due to mechanical loading exceeding the altered load-bearing threshold level.

We acknowledge that limitations exist within our study design. First, rats are weight bearing quadrupeds throughout their lifetime, while human infants outgrow crawling. The different relative time-periods of walking on forelimbs and hindlimbs between rats and humans may induce differing mechanical loading. However, neither rats nor humans use their affected arm following injury, and the murine model has been proven to produce consequences similar to humans in terms of severity (Li 2010).^(27)^

In addition, this study was not powered to detect the small differences measured in trabecular TMD and Tb.Th, but a post-hoc power analysis revealed that the measure differences would require a sample size of 16 per group for TMD and 14 per group for Tb.Th to reach statistical significance (p < 0.05). While micro-CT analyses reveal trabecular bone changes following neurectomy, the mechanisms underlying these changes are not well understood. Histological assessments of bone remodeling may provide additional insight into the underlying mechanisms driving the trabecular bone changes with peripheral nerve injury.

Finally, nerve injury is complex. Whether nerve injury directly influences skeletal development remains an open question, one that is also confounded by cellular crosstalk between muscle and bone cells (Hamrick, McNeil, & Patterson, 2010).^(28)^ A recent study reported that location of nerve injury with respect to the dorsal root ganglion can have different effects on muscle spindle preservation and longitudinal muscle growth (Nikolaou *et al.*, 2015),^(18)^ which motivates the need to examine these effects in more detail. Moreover, almost nothing is known about the timing of nerve injury during postnatal development on the serial progression of muscle and bone deficits following injury, which may provide essential information that could lead to improved therapies in human patients.

Our study reveals significantly altered trabecular bone properties in regions of the shoulder underlying the glenohumeral joint surfaces, particularly in the scapula. While fracture more commonly occurs in the clavicle as opposed to the glenohumeral bones (Ogden, 2000),^(29)^ following BPBI, our study suggests that altered trabecular bone mineralization and microstructure in the glenohumeral bones may compromise bone strength during the critical period of postnatal skeletal development. Longitudinal studies are necessary to assess whether increased risk of fracture is an overlooked, unmet clinical need. Shoulder joint health in BPBI patients may benefit from the development of better post-injury care plans that target the maintenance of underlying trabecular bone. Our study is the first to characterize trabecular microstructure in the humeral metaphyseal and the glenoid fossa region of the scapula, giving insight into the underlying microstructure of the glenohumeral as a whole post-injury.

## References

1. Natural_history_of_obstetric_b.pdf.

2. Al-Qattan MM. Classification of Secondary Shoulder Deformities in Obstetric Brachial Plexus Palsy. J. Hand Surg. 2003 Oct;28(5):483–6.

3. Nikolaou, S, Peterson, E, Kim, A, Wylie, C, Cornwall R. Impaired Growth of Denervated Muscle Contributes to Contracture Formation Following Neonatal Brachial Plexus Injury: J. Bone Jt. Surg. 2011 Mar;93(5):461–70.

4. Crouch, DL, Plate, JF, Li, Z, Saul KR.. Computational Sensitivity Analysis to Identify Muscles That Can Mechanically Contribute to Shoulder Deformity Following Brachial Plexus Birth Palsy. J. Hand Surg. 2014 Feb;39(2):303–11.

5. Crouch, DL, Hutchinson, ID, Plate, JF, Antoniono, J, Gong, H, Cao, G, Li, Z, Saul KR. Biomechanical Basis of Shoulder Osseous Deformity and Contracture in a Rat Model of Brachial Plexus Birth Palsy: J. Bone Jt. Surg. 2015 Aug;97(15):1264–71.

6. Cheng, W, Cornwall, R, Crouch, DL, Li, Z, Saul KR. Contributions of Muscle Imbalance and Impaired Growth to Postural and Osseous Shoulder Deformity Following Brachial Plexus Birth Palsy: A Computational Simulation Analysis. J. Hand Surg. 2015 Jun;40(6):1170–6.

7. Vico, L, Collet, P, Guignandon, A, Lafage-Proust M-H, Thomas, T, Rehailia, M, Alexandre C. Effects of long-term microgravity exposure on cancellous and cortical weight-bearing bones of cosmonauts. The Lancet. 2000 May;355(9215):1607–11.

8. Kazakia, GJ, Tjong, W, Nirody, JA, Burghardt, AJ, Carballido-Gamio, J, Patsch, JM, Link, T, Feeley, BT, Benjamin Ma C.. The influence of disuse on bone microstructure and mechanics assessed by HR-pQCT. Bone. 2014 Jun;63:132–40.

9. Eser, P, Schiessl, H, Willnecker J. Bone loss and steady state after spinal cord injury: A cross-sectional study using pQCT.:3.

10. Kim, HM, Galatz, LM, Patel, N, Das, R, Thomopoulos S. Recovery Potential After Postnatal Shoulder Paralysis: An Animal Model of Neonatal Brachial Plexus Palsy. J. Bone Jt. Surg.- Am. Vol. 2009 Apr;91(4):879–91.

11. Kim, HM, Galatz, LM, Das, R, Patel, N, Thomopoulos S. Musculoskeletal deformities secondary to neurotomy of the superior trunk of the brachial plexus in neonatal mice. J. Orthop. Res. 2010 Mar 11;28(10):1391–8.

12. Potter, R, Havlioglu, N, Thomopoulos S. The developing shoulder has a limited capacity to recover after a short duration of neonatal paralysis. J. Biomech. 2014 Jul;47(10):2314–20.

13. [Frost 1994] wolf’s law.pdf.

14. Turner CH. Three rules for bone adaptation to mechanical stimuli. Bone. 1998 Nov;23(5):399–407.

15. Reading, BD, Laor, T, Salisbury, SR, Lippert, WC, Cornwall R. Quantification of Humeral Head Deformity Following Neonatal Brachial Plexus Palsy: J. Bone Jt. Surg.-Am. Vol. 2012 Sep;94(18):e136-1–8.

16. Waters, PM, Smith, GR, Jaramillo D. Glenohumeral Deformity Secondary to Brachial Plexus Birth Palsy: J. Bone Jt. Surg. 1998 May;80(5):668–77.

17. Li, Z, Ma, J, Apel, P, Carlson, CS, Smith, TL, Koman LA. Brachial Plexus Birth Palsy– Associated Shoulder Deformity: A Rat Model Study. J. Hand Surg. 2008 Mar;33(3):308–12.

18. Nikolaou, S, Hu, L, Cornwall R. Afferent Innervation, Muscle Spindles, and Contractures Following Neonatal Brachial Plexus Injury in a Mouse Model. J. Hand Surg. 2015 Oct;40(10):2007–16.

19. Crouch, DL, Hutchinson, ID, Plate, JF, Antoniono, J, Gong, H, Cao, G, Li, Z, Saul KR. Biomechanical Basis of Shoulder Osseous Deformity and Contracture in a Rat Model of Brachial Plexus Birth Palsy: J. Bone Jt. Surg.-Am. Vol. 2015 Aug;97(15):1264–71.

20. Kothary, S, Rosenberg, ZS, Poncinelli, LL, Kwong S. Skeletal development of the glenoid and glenoid–coracoid interface in the pediatric population: MRI features. Skeletal Radiol. 2014 Sep;43(9):1281–8.

21. Hildebrand, T, Rüegsegger P. Quantification of Bone Microarchitecture with the Structure Model Index. Comput. Methods Biomech. Biomed. Engin. 1997 Jan;1(1):15–23.

22. Bouxsein, ML, Boyd, SK, Christiansen, BA, Guldberg, RE, Jepsen, KJ, Müller R. Guidelines for assessment of bone microstructure in rodents using micro-computed tomography. J. Bone Miner. Res. 2010 Jun 7;25(7):1468–86.

23. The compressive behavior of bone as a two-phase porous structure.pdf.

24. Parfitt, AM, Mathews, CH, Villanueva, AR, Kleerekoper, M, Frame, B, Rao DS. Relationships between surface, volume, and thickness of iliac trabecular bone in aging and in osteoporosis. Implications for the microanatomic and cellular mechanisms of bone loss. J. Clin. Invest. 1983 Oct;72(4):1396–409.

25. Stein, EM, Kepley, A, Walker, M, Nickolas, TL, Nishiyama, K, Zhou, B, Liu, XS, McMahon, DJ, Zhang, C, Boutroy, S, Cosman, F, Nieves, J, Guo, XE, Shane E. Skeletal Structure in Postmenopausal Women With Osteopenia and Fractures Is Characterized by Abnormal Trabecular Plates and Cortical Thinning: SKELETAL STRUCTURE IN OSTEOPENIC POSTMENOPAUSAL WOMEN. J. Bone Miner. Res. 2014 May;29(5):1101–9.

26. Legrand, E, Chappard, D, Pascaretti, C, Duquenne, M, Krebs, S, Rohmer, V, Basle M-F, Audran M. Trabecular bone microarchitecture, bone mineral density, and vertebral fractures in male osteoporosis. J. Bone Miner. Res. 2000;15(1):13–19.

27. Li, Z, Barnwell, J, Tan, J, Koman, LA, Smith BP. Microcomputed Tomography Characterization of Shoulder Osseous Deformity After Brachial Plexus Birth Palsy: A Rat Model Study: J. Bone Jt. Surg.-Am. Vol. 2010 Nov;92(15):2583–8.

28. Hamrick, MW, McNeil, PL, Patterson SL. Role of muscle-derived growth factors in bone formation. 2013;12.

29. John A. Ogden. Skeletal injury in the child. 3rd ed. New York: Springer-Verlag New York; 2000.

